# Population ecology of seabirds in Mexican Islands at the California Current System

**DOI:** 10.1101/2021.10.04.463095

**Authors:** Federico Méndez Sánchez, Yuliana Bedolla Guzmán, Evaristo Rojas Mayoral, Alfonso Aguirre-Muñoz, Patricia Koleff, Alejandro Aguilar Vargas, Fernando Álvarez Santana, Gustavo Arnaud, Alicia Aztorga Ornelas, Luis Felipe Beltrán Morales, Maritza Bello Yáñez, Humberto Berlanga García, Esmeralda Bravo Hernández, Ana Cárdenas Tapia, Aradit Castellanos Vera, Miguel Corrales Sauceda, Ariana Duarte Canizales, Alejandra Fabila Blanco, María Félix Lizárraga, Anely Fernández Robledo, Julio Hernández Montoya, Alfonso Hernández Ríos, Eduardo Iñigo Elías, Ángel Méndez Rosas, Braulio Rojas Mayoral, Fernando Solís Carlos, Alfredo Ortega-Rubio

## Abstract

The Baja California Pacific Islands (BCPI) is a seabird hotspot in the southern California Current System supporting 129 seabird breeding populations of 23 species and over one million birds annually. These islands had a history of environmental degradation because of invasive alien species, human disturbance, and contaminants that caused the extirpation of 27 seabird populations. Most of the invasive mammals have been eradicated and breeding colonies have been restored with social attraction techniques. We have systematic information for most of the breeding populations since 2008. To assess population trends, we analyzed data and present results for 19 seabird species on ten island groups. The maximum number of breeding pairs for each nesting season was used to estimate the population growth rate (λ) for each species at every island colony. We performed a nonparametric bootstrapping to assess whether seabird breeding populations are increasing or decreasing. San Benito, Natividad, and San Jerónimo are the top three islands in terms of abundance of breeding pairs. The most widespread species is Cassin’s Auklet with 14 colonies. Twenty-three populations of 13 species are significantly increasing while eight populations of six species are decreasing. We did not find statistical significance for 30 populations, however, 20 have λ>1 which suggest they are growing. Seven of the 18 species for which we estimated a regional population trend are significantly increasing, including three surface-nesting species: Brown Pelican, Elegant Tern and Laysan Albatross, and four burrow-nesting species: Ainley’s and Ashy Storm-Petrels, and Craveri’s and Guadalupe Murrelet. Our results suggest that the BCPI support healthy and growing populations of seabirds that have shown to be resilient to extreme environmental conditions such as the “Blob”, and that such resilience has been strengthen from conservation and restoration actions such as the eradication of invasive mammals and social attraction techniques.

## Introduction

Of all the birds in the world, seabirds are the most threatened group and the one with the greatest and fastest declines (Oro & Martínez-Abraín 2009, Croxall et al. 2012, Paleczny et al. 2015, Votier & Sherley 2017). Invasive alien species are the prevalent threat to seabirds, followed by fisheries bycatch, climate change and severe weather, pollution, and human disturbance (Croxall et al. 2012, Dias et al. 2019). North American seas and islands support nearly half of all seabird species globally (Croxall et al. 2012, NABCI 2016), mostly due to the high productivity associated with the California Current System (CCS; Oedekoven et al. 2001, Sydeman et al. 2009, Ainley & Hyrenbach 2010, Nur et al. 2011). This highly productive upwelling ecosystem is a hotspot for seabirds, where their abundance and population trends have been long and well-studied in the Canadian and United States portions of the CCS (Mason et al. 2007, Gaston et al. 2009, Ainley & Hyrenbach 2010, Nur et al. 2011) but not further south in Mexico in a comprehensive and systematic manner. For the Mexican portion of the CCS there is no comprehensive assessment of at-sea seabird abundances while the last regional multispecies population estimates for breeding colonies in the Baja California Pacific Islands (BCPI; from Coronado Archipelago in the north to the Magdalena Bay islands in the south) are from the period 1999-2003, when the BCPI harbored half (2,433,000) of the breeding individuals and 22 out of 37 taxa that occurred in the whole CCS (Wolf et al. 2006).

Mexico is home to a third of the world’s 368 seabird species (Croxall et al. 2012, Dias et al. 2019). Twenty (16%) of the 126 seabird species in Mexico are threatened: three species as Critically Endangered, four as Endangered, and thirteen as Vulnerable on the IUCN Red List of Threatened Species (IUCN 2021), while 24 (19%) are federally listed in Mexico’s Official Norm for species at-risk (*NOM-059-SEMARNAT-2010*; Berlanga et al. 2008, SEMARNAT 2021). In terms of endemism (i.e., endemic and semi-endemic species; González García & Gómez de Silva 2002), Mexico is the second most important country—just behind New Zealand—with 12 species: Craveri’s Murrelet (*Synthliboramphus craveri*), Guadalupe Murrelet (*S. hypoleucus*), Ainley’s Storm-Petrel (*Hydrobates cheimomnestes*), Black Storm-Petrel (*H. melania*), Guadalupe Storm-Petrel (*H. macrodactylus*), Least Storm-Petrel (*H. microsoma*), Townsend’s Storm-Petrel (*H. socorroensis*), Elegant Tern (*Thalasseus elegans*), Heermann’s Gull (*Larus heermanni*), Yellow-footed Gull (*L. livens*), Black-vented Shearwater (*Puffinus opisthomelas*), and Townsend’s Shearwater (*P. auricularis*) (Croxall et al. 2012, SEMARNAT 2021).

Most of the seabird populations on the BCPI were severely reduced during the 20th century (Wolf 2006, Aguirre-Muñoz et al. 2018), with 27 seabird populations extirpated due to the combined negative effects of invasive alien mammals and direct human disturbance (McChesney & Tershy 1998, Wolf et al. 2006, Bedolla-Guzmán et al. 2019). Much has changed for seabird conservation in Mexico during the last couple of decades, particularly in the southernmost region of the CCS (i.e., the Pacific Ocean off the Baja California Peninsula). From an almost complete lack of knowledge and inaction by the late 90’s, where little was known about seabird populations (Everett & Anderson 1991, McChesney & Tershy 1998) and no protection of their colonies existed (Wolf et al. 2006, Aguirre-Muñoz et al. 2008), Mexico has taken bold conservation actions, including the legal protection of its nearly 4,500 islands (Aguirre-Muñoz & Méndez-Sánchez 2017), the removal of invasive mammals from 39 islands (Aguirre-Muñoz et al. 2011, 2018), the restoration and long-term monitoring of seabird populations (Hernández-Montoya et al. 2014, Bedolla-Guzmán et al. 2017, 2019), and the formulation of a National Action Program for Seabird Conservation (*PACE Aves Marinas*; Méndez Sánchez et al. In prep., SEMARNAT 2021).

Benefits derived from the eradication of invasive mammals on the biodiversity of the world’s islands have been greatly documented (Graham et al. 2018, Jones et al. 2016, Towns et al. 2011, 2013), and many studies have focused on the recovery of seabird populations (e.g., Buxton et al. 2014, Borrelle et al. 2016, Brooke et al. 2017). On a global assessment, Brooke et al. (2017) found that after successful invasive mammal eradication, the median growth rate for 181 seabird populations of 69 species was λ=1.119, and populations with positive growth greatly outnumbered those in decline (λ > 1; n= 151 *vs* λ < 1; n = 23). Mexico is among the countries that most invasive mammal eradications has successfully implemented with 60 populations from 39 islands (Aguirre-Muñoz et al. 2011, 2018, Jones et al. 2016), and some of the few that actively conduct pre- and post-eradication monitoring (Aguirre-Muñoz et al. 2016, Towns 2018) to assess ecological outcomes (Aguirre-Muñoz et al. 2011, 2018, Bedolla-Guzmán et al. 2019, Hernández-Montoya et al. 2014, Luna-Mendoza et al. 2019, Ortiz-Alcaraz at al. 2016, 2019, Samaniego-Herrera et al. 2011, Samaniego-Herrera & Bedolla-Guzmán 2012) as well as active restoration such as seabird social attraction (Jones & Kress 2012, Kappes & Jones 2014, Pacific Rim Conservation et al. 2021) once invasive mammals are removed to maximize conservation gains (Aguirre-Muñoz et al. 2011, 2018, Bedolla-Guzmán et al. 2019). Benefits to seabird populations in Mexico from both passive and active restoration have also been documented, with the encouraging outcome that to date 23 out of 27 (85%) historically extirpated seabird breeding colonies have been restored, and 12 new breeding colonies have been recorded (Bedolla-Guzmán et al. 2019).

Beyond updating the number of breeding individuals at their island colonies (Wolf et al. 2006, Hernández-Montoya et al. 2014, Albores-Barajas et al. 2018, Whitworth et al. 2018, 2020, 2021, Bedolla-Guzmán et al. 2019), there have not been any attempts to understand the dynamics of the BCPI seabird populations and to assess recovery or decline trends at the subpopulation and metapopulation level (Buckley & Downer 1992, Munilla et al. 2016). Therefore, building upon our long-term monitoring of the seabird populations on the BCPI for the past 17 years, **in this paper we aim to answer two questions: (1) What is the breeding status and spatial diversity patterns of the seabird populations on the BCPI?; and (2) What are their population trends: growing, declining or no change?** By doing so, we expect to highlight the importance and the contributions of these populations and their habitats—islands and the surrounding marine environment within the CCS—in the context of seabird conservation in the central eastern Pacific and the world.

## Methods

### Study area

The Baja California Pacific Islands (BCPI) are in the southern California Current System, off the Baja California Peninsula (Fig 1). In this region, there are around 30 islands, all within Protected Areas managed by Mexico’s Federal Government through the National Commission for Natural Protected Areas (CONANP): El Vizcaíno Biosphere Reserve; Guadalupe Island Biosphere Reserve; and Baja California Pacific Islands Biosphere Reserve (Aguirre-Muñoz & Méndez-Sánchez 2017, UNEP-WCMC 2020). They are key reproduction sites for 133 species of vertebrates: 41 amphibians and reptiles, 69 birds, 19 mammals and four pinnipeds (Latofski-Robles et al. 2014). We focus our analyses on the ten islands and archipelagos described in Table 1. Due to their relevance for birds, all these islands are Important Bird and Biodiversity Areas (IBAs; Arizmendi & Vázquez Valdelamar 2000, Vidal et al. 2009). The size of the islands where we conducted our work ranges from 35 to 24,171 hectares, with altitudes ranging from 10 to 1,298 meters. Except for San Martín and Guadalupe islands, which have a volcanic origin, the rest are an unsubmerged extension of the continental shelf (Samaniego Herrera et al. 2007). Most of them are within proximity to the Baja California Peninsula, between 1.8 to 72 km, with Guadalupe Island being the most oceanic at 260 km. This region is characterized by a Mediterranean climate, with hot and dry summers and cold and wet winters, a regional average annual temperature of 18-23°C, and an average annual cumulative precipitation of *ca*. 200 millimeters. The dominant plant communities are maritime desert scrub, although Guadalupe Island sustains a temperate forest because of its high elevation and an almost permanent fog system (Luna-Mendoza et al. 2019).

**Fig 1.**
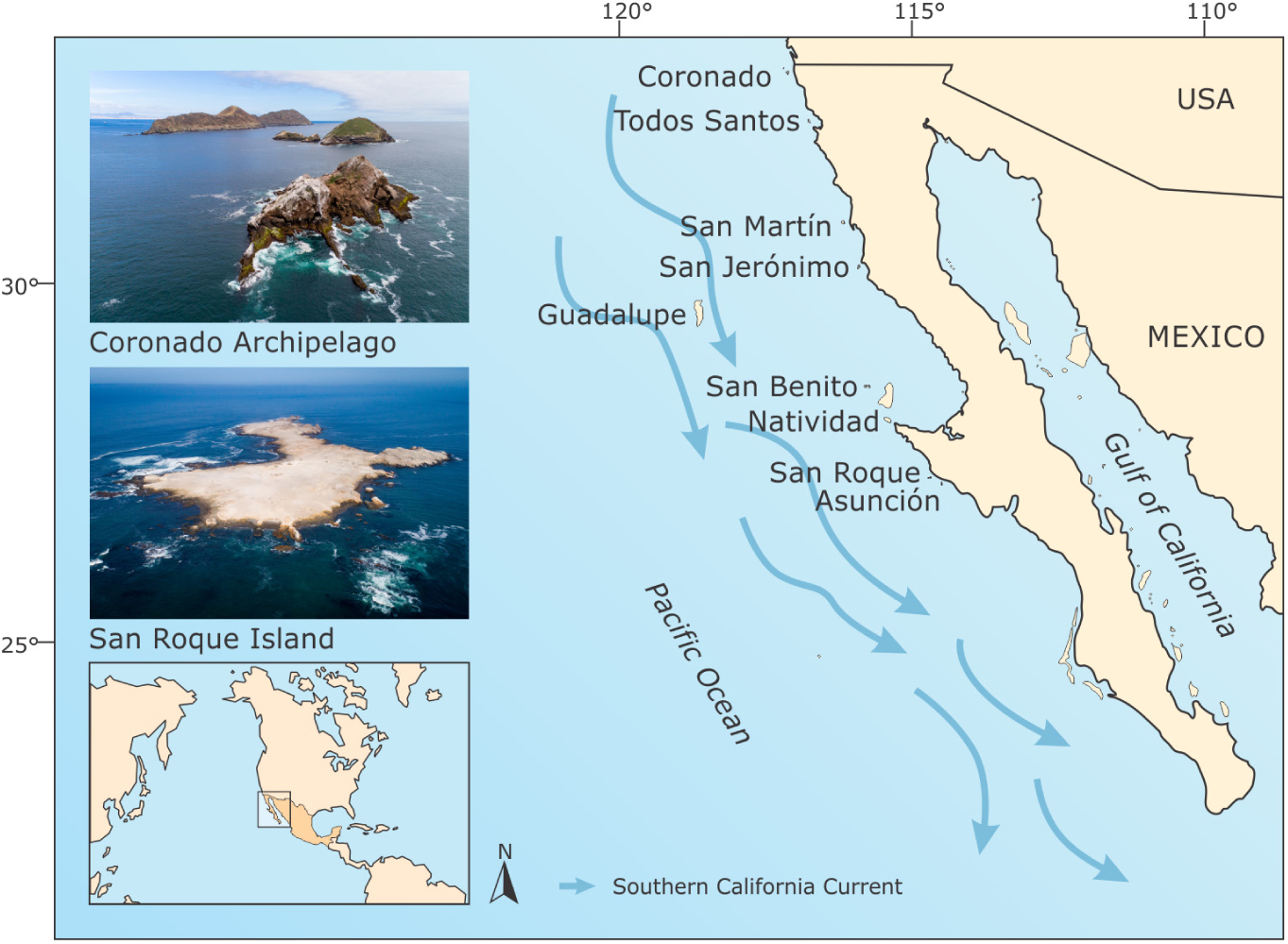
Map of the Baja California Pacific Islands, a seabird hotspot where breeding populations have been systematically monitored for almost two decades. Photos of Coronado Archipelago and San Roque islands are shown—being the extremes in geographic location—to show the heterogenous physiography of the region’s islands. Arrows depict the southerly flow of the California Current (Sydeman & Elliott 2008), which has significant influence on the region’s marine productivity and thus the seabird populations. Photo credits: © GECI / J.A. Soriano.

**Table 1.** Characteristics of the Baja California Pacific Islands where breeding seabird populations were monitored during the period 2003-2019.

| Island/Archipelago | Area (Hectares) | Protected Area<sup>a</sup> | No. breeding seabird species<sup>b</sup> |
| --- | --- | --- | --- |
| Coronado (3 islands, 1 islet) | 173 | PIBR | 11 |
| Todos Santos (2 islands) | 123 | PIBR | 8 |
| San Martín | 265 | PIBR | 10 |
| San Jerónimo | 48 | PIBR | 11 |
| Guadalupe | 24,171 | GIBR | 5 |
| Morro Prieto and Zapato (2 islets) | 45 | GIBR | 8<sup>c</sup> |
| San Benito (3 islands) | 610 | PIBR | 13 |
| Natividad | 736 | EVBR | 7 |
| San Roque | 35 | EVBR | 9 |
| Asunción | 43 | EVBR | 7 |

<sup>a</sup>PIBR: Baja California Pacific Islands Biosphere Reserve; GIBR: Guadalupe Island Biosphere Reserve; EVBR: El Vizcaíno Biosphere Reserve. <sup>b</sup>Updated from: Wolf et al. (2006), Whitworth et al. (2018, 2020, 2021), Bedolla-Guzmán et al. (2019) and present work. <sup>c</sup>Morro Prieto and Zapato host 6 and 7 species each; collectively, they harbor 8 species (5 shared between them).

### Monitoring of seabird colonies

We have been monitoring seabird populations on Guadalupe Island since 2003 (Hernández-Montoya et al. 2014, 2019), on Asunción and San Roque islands since 2008 (Aguirre-Muñoz et al. 2011, Bedolla-Guzmán et al. 2019), and from 2014 expanded to the rest of the islands in the region (except Magdalena Bay islands; Aguirre-Muñoz et al. 2018, Bedolla-Guzmán et al. 2019). For this study, our data sample includes 61 breeding colonies of 19 seabird species: 5 Procellariiformes, 9 Charadriiformes, 4 Suliformes, and 1 pelican, on ten islands and archipelagos (S1 Table).

We used the maximum number of breeding pairs for each nesting season to estimate the population growth rate (λ, lamda; S2 Table). For surface nesting-species (i.e., cormorants, pelicans, terns, and gulls), we surveyed active nests from land-based vantage points, complemented with surveys around the islands (boat counts), every 15 days during the whole breeding season. For burrow-nesting species (i.e., shearwaters, petrels, auklets, and murrelets), we conducted an exhaustive and intensive search of active burrows in all potential breeding sites. Burrow occupancy was determined either by recording apparently occupied burrows (i.e., with signs of activity such as guano, feathers, clear entrances, and footprints; Walsh, et al. 1995) or by directly confirming burrow content (i.e., adult, egg, or chick). On islands with accessible nesting sites (i.e., Asunción and San Roque), we conducted a census across the whole island, and either recorded apparently occupied burrows or checked burrow content using a hand-lamp or a burrowscope. Pairs nesting on artificial colonies installed on all the islands for social attraction (Bedolla-Guzmán et al. 2019) were included on our counts. For those species with high nest density such as the Western Gull (*Larus occidentalis*) on the Todos Santos Archipelago, Cassin’s auklet (*Ptychoramphus aleuticus*) on San Jerónimo Island, and Black-vented Shearwater (*Puffinus opisthomelas*) on Natividad Island, we estimated nest or burrow densities during peak incubation, counting nests or burrows within circular or square plots randomly distributed and georeferenced (Keitt et al. 2003, Parker & Rexer-Huber 2020). Bootstrap percentile intervals for the population size were calculated by means of bootstrapping (Berrar 2019) using the sampling plots of nests and occupied burrows.

### Spatial diversity patterns of seabirds

We analyzed spatial diversity patterns of seabirds on the BCPI through diversity indices (Koleff et al. 2003, Begon et al. 2006) using all available data shown in S2 Table. We calculated values of indices for abundance (i.e., number of breeding pairs) and for presence-absence data to evaluate the geographical structure of the seabird populations. For alpha diversity we used the Shannon Index (*H′*) and the inverse Simpson Index 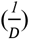 as per equations (1) and (2), respectively, where *S* is the total number of species and *p*_*i*_ is the proportional abundance for each species on a given island or archipelago.

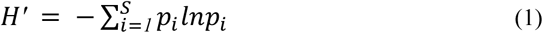

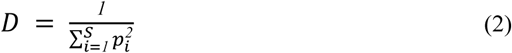

For beta diversity (*β*_*t*_) we calculated the Sørensen Index of dissimilarity (Wilson & Shmida 1984, Koleff et al. 2003) using *vegan: Community Ecology Package* for R version 2.5-6 (Oksanen et al. 2019) that uses equation (3), where *a* is the total number of species on two islands (one of interest and one of reference); *b* is the amount of species present on the reference island that are not present on the interest island, while *c* is the opposite.

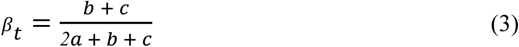

### Population growth trends

#### Model

We estimated the annual population growth rates (λ) for each species nesting on each island or archipelago using the relationship described in equation (4):

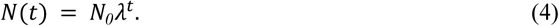

*N*_*0*_is the number of breeding pairs at the time when the first count of the period was made, λ is the annual population growth rate and *t* is the time interval between the first and last counts of the period. This was done for 19 seabird species (S2 Table). With results from equation (4) we calculated the percent of change in the population using equation (5):

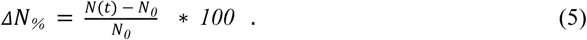

We calculated a regional population growth rate (λ_R_) and its percent of change using equations (4) and (5), respectively. We did this for those species with at least three consecutive years of data on all their breeding colonies in the different islands (18 of the 19 monitored species; Royal Tern was not included). We used the sum of the breeding numbers for all colonies for common years as the number of breeding pairs at the time *t*. For instance, for Brandt’s Cormorant we only have data for all its breeding colonies in the period 2016-2018, although we have monitored some of its colonies since 2012 (e.g., Asunción and San Roque).

#### Nonparametric Bootstrapping

We performed a nonparametric bootstrapping (Berrar 2019) to assess whether seabird breeding populations are increasing, decreasing or have no significant change (i.e., undetermined). Nonparametric bootstrapping makes no assumptions on data distribution and sampling is done with replacement (Berrar 2019). We generated a bootstrap set *N*^*^ by randomly sampling *n* instances with replacement from our data set of maximum number of breeding pairs (S2 Table), *N*_*obs*_, where *n* is the total number of records (i.e., monitored breedings seasons; e.g., n= 5 for Brandt’s Cormorant on Natividad Island). The sampling is uniform, meaning each of the *n* elements in *N*_*obs*_ has the same probability 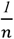 of being selected (Berrar 2019). We then calculated the bootstrap version *λ*\*, of the population growth rate *λ*, by fitting equation (1) to the bootstrap sample *N*^*^. We repeated these last steps *b* = *2000* times and removed the outliers. We defined outliers as the values for each *λ*\* distribution outside the interval [*Q*_*1*_ − *k* ⋅ *IQR, Q*_*3*_ + *k* ⋅ *IQR*]. This is the Tukey fences criterion, where *Q*_*1*_ is the first quartile, *Q*_*3*_ the third quartile, *IQR*= *Q*_*3*_ – *Q*_*1*_ is the interquartile range, and *k* is a positive constant that defines the outlier tolerance level. Here we used *k* = *1*.*5* as is customary. After this, we calculated the 95% bootstrap percentile interval for *λ*\*. Finally, we tested the following null hypotheses: increasing population, *H*_*0*_: *λ* ≤ *1, p* < *α* = *0*.*1* and decreasing population, *H*_*0*_: *λ* ≥ *1, p* < *α* = *0*.*1*.

## Results

### Status of the seabird breeding populations

The BCPI current seabird assemblage is 23 species and a total of 129 breeding colonies on 17 islands. Table 2 contains the most recent and updated information to date on the number of breeding pairs per seabird species per island on the BCPI. On average for the period 2014-2019, the BCPI supported 323,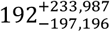 breeding pairs at 61 colonies of 19 species (48% and 83% of the total, respectively). This means that considering just half the colonies (61 vs 126), at least between ca. 251,000 and 1.1 million individuals breed on this important seabird hotspot every year. This figure is like the estimates by Mason et al. (2007) from aerial at-sea and coastal surveys for seabirds off southern California (from Cambria, California, USA, to the Mexican border) in 1999-2002, and below the estimate of 2.4 million breeding pairs for the BCPI region done by Wolf et al. (2006) from a literature review and censuses in 1999-2003. San Benito, Natividad, and San Jerónimo are the top three islands in terms of abundance of breeding pairs (2014-2019 average): 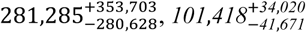 and 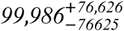, respectively.

**Table 2.** Breeding status of the seabird populations on the Baja California Pacific Islands for the 2017-2019 breeding seasons.

| Name | Species | Coronado Norte | Coronado Sur | Terrón de Azúcar | Coronado Centro | Todos Santos Sur | Todos Santos Norte | San Martín | San Jerónimo | Guadalupe | Zapato | Morro Prieto | San Benito Oeste | San Benito Medio | San Benito Este | Natividad | San Roque | Asunción |
| --- | --- | --- | --- | --- | --- | --- | --- | --- | --- | --- | --- | --- | --- | --- | --- | --- | --- | --- |
| Laysan Albatross | *Phoebastria immutabilis* |  |  |  |  |  |  |  |  | 395<sup>(e)</sup> | 745<sup>(e)</sup> | 371<sup>(e)</sup> |  |  |  |  |  |  |
| Black-vented Shearwater | *Puffinus opisthomelas* |  |  |  |  |  |  |  |  | E | 229 | 400 (+120 -110) | 86 | 11 | 24 | 118,920 (+16,740 -16,290) |  |  |
| Leach's Storm-petrel | *Hydrobates leucorhous* | E |  | 2 |  |  |  |  |  |  |  |  | B | B | B |  |  |  |
| Townsend's Storm-petrel | *Hydrobates socorroensis* |  |  |  |  |  |  |  |  | 2 | 11 | 890 (+520 -420) |  |  |  |  |  |  |
| Ainley's Storm-petrel | *Hydrobates cheimomnestes* |  |  |  |  |  |  |  |  | PB | 2<sup>(e)</sup> | 181<sup>(e)</sup> |  |  |  |  |  |  |
| Ashy Storm-petrel | *Hydrobates homochroa* | 5 |  |  |  | 20 | 55 | PB |  |  |  |  |  |  |  |  |  |  |
| Black Storm-petrel | *Hydrobates melania* | 104<sup>(a)</sup> | PB | 52<sup>(a)</sup> |  |  |  | PB |  |  |  |  | B | B | B |  |  |  |
| Least Storm-petrel | *Hydrobates microsoma* |  |  |  |  |  |  |  |  |  |  |  | B | B | B |  |  |  |
| Brown Pelican | *Pelecanus occidentalis* | 597 | 442 |  |  | 723 | E | 376 | 166 |  |  |  |  |  | 78 | 531 | 37 | 243 |
| Blue-footed Booby | *Sula nebouxii* |  |  |  |  |  |  |  | 1 |  |  |  |  |  |  |  |  | 2(g) |
| Brown Booby | *Sula leucogaster* |  |  | 13 |  |  |  |  |  |  | 27 (+8 -8) |  |  |  |  |  |  |  |
| Double-crested Cormorant | *Phalacrocorax auritus* | 56 | 164 |  |  | 238 | E | 791 | 104 |  |  |  |  |  | 39 | 823 | 113 | 131 |
| Brandt's Cormorant | *Phalacrocorax penicillatus* | 52 | B | 2 | 2 | 566 | B | 130 | 180 |  | 39 (+5 -5) |  |  |  | 22 | 2,400 | 4,747 | 3,228 |
| Pelagic Cormorant | *Phalacrocorax pelagicus* | 1 | 3 | 5 |  | 19 | 2 |  | 1 |  |  |  |  |  |  |  |  |  |
| Heermann's Gull | *Larus heermanni* |  |  |  |  |  |  |  |  |  |  |  |  | 121 |  |  | 42 |  |
| Western Gull | *Larus occidentalis* |  | 194 |  |  | 5,990<br>(+1,100<br>-1,130) | 2,470<br>(+820<br>-740) | 1,382 | 2,442 | B |  |  | B | 568 | 442 | 300<sup>(f)</sup> | 1,749 | 1,323 |
| Caspian Tern | *Hydroprogne caspia* |  |  |  |  |  |  | 186 | 18 |  |  |  |  |  |  |  |  |  |
| Elegant Tern | *Thalasseus elegans* |  |  |  |  |  |  |  | 684 |  |  |  |  |  |  |  | 2,500 | E |
| Royal Tern | *Thalasseus maximus* |  |  |  |  |  |  |  | 171 |  |  |  |  |  |  |  | 870 |  |
| Scripps's Murrelet | *Synthliboramphus scrippsi* | 25<sup>(a)</sup> | 34<sup>(a)</sup> | 27<sup>(a)</sup> | 11<sup>(a)</sup> | 84 | 17 | B | 3 |  |  |  | 145 | 7 | 18 |  |  |  |
| Guadalupe Murrelet | *Synthliboramphus hypoleucus* |  |  |  |  |  |  |  |  | 275<sup>(e)</sup> | 2,360<br>(+530<br>-500) | 1,600<br>(+480<br>-420) | 2 | 1 |  |  |  |  |
| Craveri's Murrelet | *Synthliboramphus craveri* |  |  |  |  |  |  | 1<sup>(b)</sup> |  |  |  |  | 1 |  |  | 1 | 5 | 1 |
| Cassin's Auklet | *Ptychoramphus aleuticus* | 1<sup>(h)</sup> | 20 | B |  | 30 | 33 | B<sup>(c)</sup> | 218,250<sup>(d)</sup><br>(+17200<br>-17410) | E |  | 202 | 472,606<br>(+62,443<br>-62,473) | 7,621 | 79,224<br>(+13,190<br>-12,541) | 12 | 1,659 | 2,602 |
| **Total breeding taxa** |  | **8** | **8** | **7** | **2** | **8** | **6** | **10** | **11** | **5** | **7** | **6** | **9** | **9** | **10** | **7** | **9** | **7** |

Records show the maximum number of breeding pairs per species per island during our 2017-2019 surveys. B: Breeder; PB: Probable breeder; E: Extirpated. <sup>(a)</sup> Maximum number recorded but no exhaustive survey was possible. <sup>(b)</sup> Record by Whitworth et al. (2018) for the 2007-2008 breeding season, during our own surveys we just found and confirmed Scripps's Murrelet. <sup>(c)</sup> 136 apparently occupied nests were found in 2017 but no exhaustive survey was possible. <sup>(d)</sup> Data from 2016. <sup>(e)</sup> Data from 2020. <sup>(f)</sup> Data from 2015. <sup>(g)</sup> New record. <sup>(h)</sup> Previously considered extirpated.

The most widespread species is Cassin’s Auklet with 14 colonies, followed by Brandt’s Cormorant with 13, Western Gull with 12 and Scripps’s Murrelet with 11. The less widespread species are Blue-footed Booby, Brown Booby, Caspian Tern, Elegant Tern, Hermann’s Gull and Royal Tern with two colonies each (S1 Appendix Fig A). This is not surprising since the three tern species and the Brown Booby are either recent records or recolonizations to the BCPI, while the Blue-footed Booby has its northernmost breeding range at San Jerónimo Island, and the Hermann’s Gull has historically been restricted to two islands with small colonies (Bedolla-Guzmán et al. 2019, Wolf et al. 2006).

The San Benito Archipelago hosts the greatest seabird assemblage with 13 breeding species, followed by the Coronado Archipelago and San Jerónimo Island (11 species), and San Martín Island (10 species) (S1 Appendix Fig B). It stands-out the number of species on San Jerónimo and San Martín despite their relatively small size (48 and 265 hectares, respectively). The San Benito Archipelago has the same number of breeding species to that reported by Wolf et al. (2006), however, the Coronado Archipelago, San Jerónimo and San Martín now host 1, 7 and 4 more breeding taxa, respectively, than two decades ago (S3 Table). These recent breeding records have been previously reported by Bedolla-Guzmán et al. (2019). We are reporting one new record, the Blue-footed Booby on Asunción Island, and one new recolonization of a previously extirpated species, the Cassin’s Auklet on Coronado Norte (S3 Table).

The island with the lowest number of breeding species is Guadalupe (5 species), despite being the biggest (24,171 ha) although the most oceanic island (S1 Appendix Figs B and C). Two species are extirpated from this island due to predation by feral cats: Black-vented Shearwater and Cassin’s Auklet, which are restricted to the nearby cat-free Zapato and Morro Prieto islets (S3 Table). There is an ongoing comprehensive restoration program on the island which includes the eradication of the cat population and the implementation of social attraction techniques to restore these extirpated populations from the islets’ colonies. This island is also the most important breeding colony in the eastern Pacific for the Laysan Albatross (Hernández-Montoya et al. 2014, 2019) and there is an ongoing conservation translocation project to restore the Black-footed Albatross (*P. nigripes*) population to mitigate the impacts of sea-level rise caused from climate change at its main breeding colonies in Hawaii (Pérez Ortega 2021).

The seabird species’ richness in the BCPI increases with island area (e.g., San Benito and San Martín) but decreases with isolation (i.e., distance from the mainland; S1 Appendix Fig C). Guadalupe Island and its islets Zapato and Morro Prieto are outliers, suggesting they should be treated separately given their oceanic origin where evolutionary dynamics vary from islands closer to the mainland (Whittaker et al. 2008). From our analyses of spatial diversity patterns, the islands that had the greatest seabird alpha diversity are San Roque, Coronado, Morro Prieto and Zapato, and San Martín, whereas the less diverse islands are Natividad and San Jerónimo. On the latter, the Shannon Index (*H′*) is low (*H′* = *0*.*103*) possibly because the dominant species is the Cassin’s Auklet despite harboring 11 species. Something similar occurs with Natividad (*H′* = *0*.*200*) where the dominant species is the Black-vented Shearwater although it harbors seven species (Table 3). With respect to beta diversity, the islands that are most similar are Natividad-Asunción, Coronado-Todos Santos, and Guadalupe-Morro Prieto/Zapato (S1 Appendix Fig D and E). This means that these islands share many species and thus have similar species composition, possibly because of their proximity.

**Table 3.** Alpha diversity of breeding seabirds on the Baja California Pacific Islands calculated from the average number of breeding pairs regardless of the season. *H′*is the Shannon Index while *D* is the Inverse Simpson Index per equations (1) and (2).

| Island/Archipelago | $H'$ | $D$ |
| --- | --- | --- |
| San Roque | 1.525 | 3.627 |
| Coronado | 1.497 | 3.648 |
| Morro Prieto and Zapato | 1.310 | 2.898 |
| San Martín | 1.307 | 3.096 |
| San Benito | 1.270 | 3.262 |
| Asunción | 1.144 | 2.622 |
| Guadalupe | 0.683 | 1.96 |
| Todos Santos | 0.639 | 1.37 |
| Natividad | 0.200 | 1.078 |
| San Jerónimo | 0.103 | 1.032 |

### Seabird population trends

We were able to assess the population growth rates of 61 colonies of 19 species of the BCPI. Estimated population growth rates and the percent of change of the populations over the monitored timeframes are shown in S1 Table. Twenty-three (37.7%) populations of 13 species are significantly increasing while eight (13.1%) populations of six species are decreasing (Fig 2 and Table 4). We did not find statistical significance for 30 (49.2%) populations thus were unable to determine an increasing nor decreasing trend. The main reason is that we do not have enough information to reject our null hypotheses. This does not mean that such populations have a stable trend but rather that we need to continue monitoring them for subsequent years to be able to find a positive or negative trend. Yet, 20 out of those 30 populations have a population growth rate of *λ* > *1*.*0*, which suggests they are indeed growing. Overall, this means that 70% (43) of the assessed seabird populations (*n* = *61*) on the BCPI show a positive growth trend (Fig 2 and S1 Table). We also found that all taxonomic groups evaluated except that of boobies and cormorants (Suliformes) are growing, and that populations of murrelets, auklets, gulls, and terns (Charadriiformes) show the fastest growth (Fig 3). The median growth rate for all seabird populations in the BCPI was λ=1.08.

**Fig 2.**
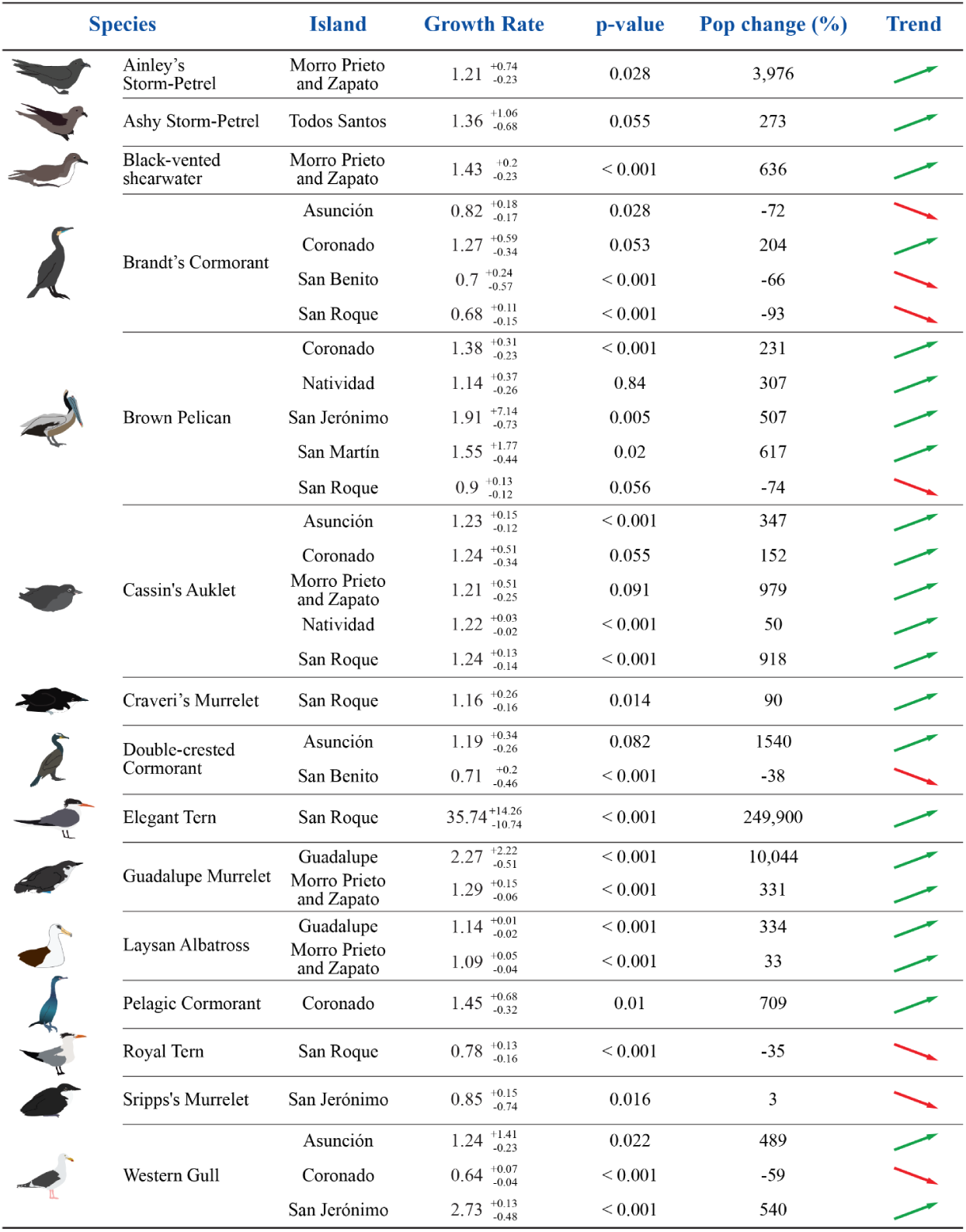
Seabird population trends on the Baja California Pacific Islands for the period 2014-2019. Only species colonies that tested the following null hypotheses: increasing population, *H*_*0*_: *λ* ≤ *1, pp* < *α* = *0*.*1* (23 colonies, 13 species) and decreasing population, *H*_*0*_: *λ* ≥ *1, pp* < *α* = *0*.*1* (8 colonies, 6 species) are shown.

**Table 4.** Summary of population trends for 61 colonies of 19 seabird species on 10 islands and archipelagos in the Mexican Pacific off the Baja California Peninsula.

| Island/Archipelago | No. Populations | Population trend status |  |  |
| --- | --- | --- | --- | --- |
|  |  | Increasing | Decreasing | Undetermined |
| Coronado | 7 | 4 | 1 | 2 |
| Todos Santos | 8 | 1 | 0 | 7 |
| San Martín | 5 | 1 | 0 | 4 |
| San Jerónimo | 8 | 2 | 1 | 5 |
| Guadalupe | 2 | 2 | 0 | 0 |
| Morro Prieto and Zapato | 6 | 5 | 0 | 1 |
| San Benito | 4 | 0 | 2 | 2 |
| Natividad | 6 | 2 | 0 | 4 |
| San Roque | 9 | 3 | 3 | 3 |
| Asunción | 6 | 3 | 1 | 2 |
| **TOTAL** | **61** | **23** | **8** | **30** |
| **%** | **100** | **38** | **13** | **49** |

**Fig 3.**
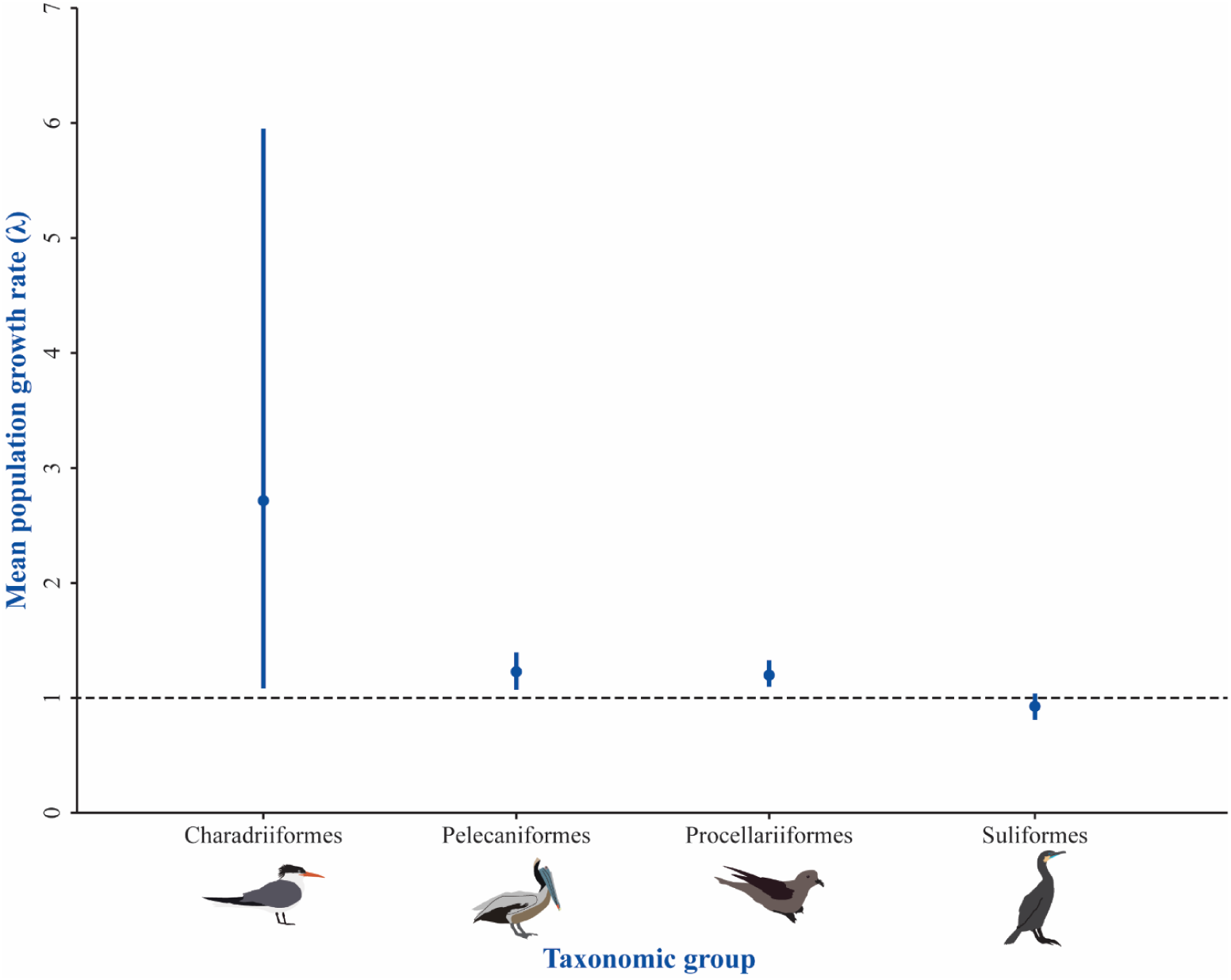
Population trends per taxonomic group. Points represent the bootstrapping means from the medians of the population growth rates (*λ*) shown in S1 Table. Bars represent the 95% bootstrap interval. The horizontal dashed line shows *λ* = *1*which indicates no population change.

Cassin’s Auklet and Brown Pelican are the species that have the most populations—five and four colonies, respectively—with a positive growing trend, while Brandt’s Cormorant has the most decreasing populations with three colonies. Three species stand out for their fast population growth rates: Elegant Tern on San Roque Island (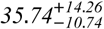; 249,900% population change 2017-2019), Western Gull on San Jerónimo Island (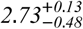; 540% population change 2015-2017), and Guadalupe Murrelet on Guadalupe Island (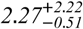; 10,044% population change 2015-2019) (Fig 2 and S1 Table). Both the Elegant Tern and Guadalupe Murrelet are newly established colonies— one pair in 2017 and two pairs in 2015, respectively—which explains its fast growth, particularly due to immigration, as found by Brooke et al. (2018).

For the period 2014-2019, seven (38.9%) of the 18 species for which we estimated a regional population trend are significantly increasing, including three surface-nesting species (Brown Pelican, Elegant Tern and Laysan Albatross) and four burrow-nesting species (Ainley’s and Ashy Storm-Petrels, and Craveri’s and Guadalupe Murrelet). Three species are restricted to the remote Guadalupe, Morro Prieto and Zapato: Ainley’s Storm-Petrel, Guadalupe Murrelet and Laysan Albatross (Fig 4). Other seven species have a population growth rate *λ* > *1*, which suggests a positive growth despite not statistically significant. This suggests that 78% (14) of the assessed species (n= 18) on the BCPI show a positive regional growth trend.

**Fig 4.**
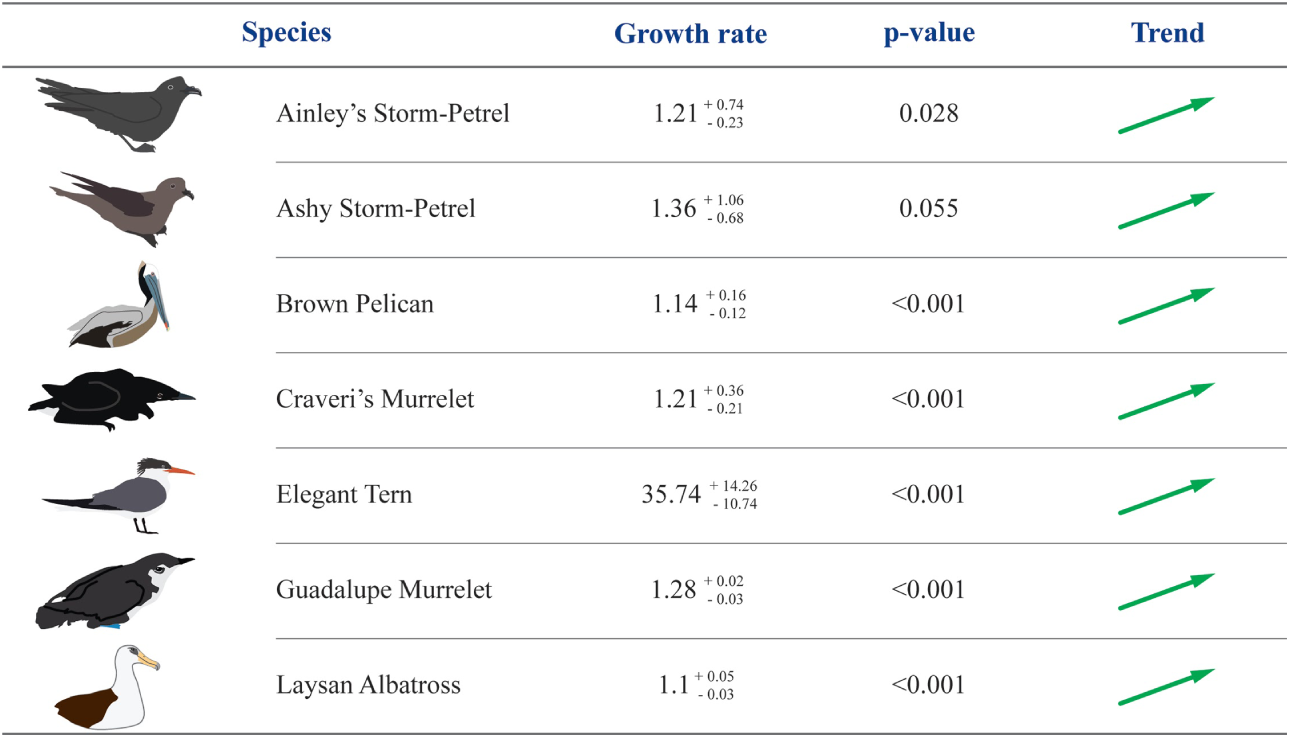
Seabirds with a positive regional population trend on the Baja California Pacific Islands for the period 2014-2019. For an increasing population, the following null hypothesis was tested: *H*_*0*_: *λ* ≤ *1, pp* < *α* = *0*.*1*.

## Discussion

Our results confirm that the BCPI are an important seabird hotspot at a national, regional, and global scale. These islands still host a good proportion of all the species distributed along the CCS and there remains to be a high breeding seabird abundance than two decades ago as reported by Wolf et al. (2006). Our estimation of ca. 251,000-1,100,000 breeding individuals is almost half than that of 2.4 million breeding birds reported by these same authors. We argue this is not because the seabird abundance has decreased in the region but more to the fact of how Wolf et al. (2006) calculated such figure from bibliographical population estimates combined with few population censuses between 1999 and 2003. Also, we consider our estimation conservative since our analyses only considered nearly half (61) of the 129 breeding colonies that currently occur within the BCPI, and we are not including breeding numbers from colonies on the islands of Magdalena Bay (i.e., Santa Magdalena, Santa Margarita and Creciente).

The San Benito Archipelago, Natividad and San Jerónimo remain to be the three islands with the greatest abundance, with the former and the latter hosting the greatest number of breeding seabird species of all the BCPI. The case of San Jerónimo stands out given that it doubled its species richness in a couple of decades, from four species reported by Wolf et al. (2006) to eleven species at present time (Table 2). This reveals the importance of this small island for the seabirds in the BCPI and thus the relevance to prevent and mitigate threats such as guano mining—still a latent danger since there exists an active lease to exploit it although it has been demonstrated that seabird guano is more valuable if retain in the ecosystem for aiding in nutrient deposition from the marine to the terrestrial environment (Plazas-Jiménez & Cianciaruso 2020)—and the introduction of invasive alien species both of which severely affected the seabird populations on this island for many decades. This increase in species might be related to the fact that the island is free from invasive mammals since 1999 (Aguirre-Muñoz et al. 2018) and that it has been protected since 2016 as part of the Baja California Pacific Islands Biosphere Reserve (Aguirre-Muñoz & Méndez-Sánchez 2017). But also, to the existence since 2014 of community-led marine reserves to protect habitat and restore populations of abalone (*Haliotis spp*.) as well as to increase fish recruitment, which have already proven to have an indirect positive effect on marine mammals such as the California Sea Lion (Zalophus californianus) (Arias-Del-Razo et al. 2019), plus the fact that San Jerónimo lies within an upwelling center making its surrounding environment highly productive (Woodson et al. 2019).

Specie richness not only remained high on the BCPI but it increased during the past couple of decades, with the number of colonies increasing twofold, from 62 (Wolf et al. 2006) to 129— partly because we surveyed all islands and islets and provide disaggregated information for each island within archipelagos in Table 3—, and with the number of species increasing from 19 (Wolf et al. 2006) to 23. San Roque, Todos Santos and Morro Prieto and Zapato are the islands that showed the highest diversity based on number of species and abundance of breeding individuals. This is the first time that diversity indices are estimated for seabirds on the BCPI thus we suggest continuing monitoring the colonies and obtaining diversity indices to evaluate spatial diversity patterns in the long term. We found three clusters of islands that share many species, two of them are close to the mainland, and in the extremes north and south: Coronado-Todos Santos (ca. 80 km apart) and Natividad-Asunción (ca. 120 km apart), while the other comprises the most offshore islands: Guadalupe-Morro Prieto/Zapato (within proximity: 1-3 km).

We provide the most comprehensive and recent breeding status of this region’s seabirds, including the first multi-colony and multi-species evaluations of population growth trends. Seabird populations in the BCPI have been showing an improvement both in number of species, colonies and breeding pairs as has been demonstrated by Bedolla-Guzmán et al. (2019) and our own present work. Yet, no previous evaluation of the trends of this seabird populations existed, particularly at a regional scale. We found that 43 out of 61 seabird populations on the BCPI are growing, which means that a third of all known breeding populations (N= 129) have a positive population growth trend. In contrast, 18 out of 61 seabird populations are declining, which represents just 14% of the whole breeding populations. The median growth rate of λ= 1.08 for all assessed seabird populations is similar to that found by Brooke et al. (2018) of λ= 1.119 for 181 seabird populations of 69 species on islands around the world. It is also similar to the population growth rate recorded by Hernández-Montoya et al. (2014), λ= 1.10, and Gallo-Reynoso & Figueroa Carranza (1996), λ= 1.35, for a steadily expanding Laysan Albatross population on Guadalupe Island. Currently, this breeding population has a λ= 1.14, just barely higher than almost a decade ago, and it remains to be higher than λ observed for other species of albatross (Weimerskirch et al. 2018). We found a high, atypical λ= 35.74 for the Elegant Tern on San Roque Island. This is explained because it is a newly established colony where the species recolonized the island in 2017 with one breeding pair, this after a decade of a systematic seabird social attraction program (Bedolla-Guzmán et al. 2019). For the following years, the colony has significantly increased in breeding numbers, which is must probably the result of immigration playing an important role in the formation of the new colony (Brooke et al. 2018). A contrasting example on this same island is that of the Royal Tern that had been extirpated for almost a century, with a λ= 0.78 despite also being a newly formed colony in the same year as that of Elegant Tern (Bedolla-Guzmán et al. 2019). For this species we suggest that the decreasing trend is mostly related to the short timeframe since the colony formation and thus would suggest keeping monitoring further years to reevaluate such trend. It might also be related to what Brooke et al. (2018) point out about growth rate declining over time because immigrant birds become a smaller fraction of the breeding population. The results obtained for this tern species can also be related to the low philopatry known for gulls and terns, where local population dynamics are influenced by emigration and immigration processes that can produce annual variations in breeding numbers (Oro & Martínez-Abraín 2009). Environmental factors have also an influence in the distribution of Elegant Tern since it has been proved that this species adapts to changing oceanographic conditions and fish availability, which makes it migrate from the Gulf of California to Southern California searching more productive waters (Velarde et al. 2015).

At a regional scale, 14 out of 23 species on the BCPI show growing populations. It stands out that three of these species occur on the Guadalupe-Morro Prieto/Zapato cluster. This is not surprising because the islets are seabird havens where no pressing threats exist and no perturbations such as the existence of invasive mammals have ever occurred, and Guadalupe Island has been subject to a comprehensive and long-term restoration program, which has had a positive effect on the Laysan Albatross population (Hernández-Montoya et al. 2019) and very recently on the Guadalupe Murrelet thanks to the protection of its potential nesting habitat from the presence of feral cats with an exclusion fence since 2014 and with the almost four-year eradication program that is being carried out on the island. The case of the Guadalupe Murrelet can be explained as a rapid population growth rate following a successful invasive mammal eradication by immigration from the nearby islets (Brooke et al. 2018).

The BCPI region has been subject to variable and extreme environmental conditions that are known to negatively affect seabird populations. The “Blob” occurred in 2013-2015 and reached the coasts of the Baja California Peninsula in May 2014, lasting until April 2015 (Bond et al. 2015). It was followed by strong ENSO conditions, “Godzilla”, that lasted until the end of 2016 (Di Lorenzo & Mantua 2016). This extreme conditions severely affected the CCS and its species, including massive die-offs of seabirds, such as Cassin’s Auklets in the central CCS in the western coast of the United States (Cavole et al. 2016). For the BCPI region, the 2014-2016 marine heatwaves have been the most intense and persistent events recorded to date (Arafeh-Dalmau et al. 2019). For instance, we recorded a 38% and 50% nest abandonment for the Brown Pelican and Brandt’s Cormorant, respectively, in 2015, between the “Blob” and “Godzilla” marine heatwaves. Nonetheless, despite these severe environmental conditions in recent years, the seabird breeding abundances and population growth rates that we report here indicate that the BCPI seabird populations are resilient to environmental variations (Velarde & Ezcurra 2018). Such resilience has and can be further strengthened from conservation actions such as the eradication of invasive mammals (Aguirre-Muñoz et al. 2011, 2018) and social attraction techniques (Bedolla-Guzmán et al. 2019).

## Supporting information

**S1 Table. Population trends for 19 seabird species at their breeding colonies on 10 islands/archipelagos in the Mexican Pacific off the Baja California Peninsula**.

A 95% credible interval was calculated for population growth rates (*λ*) using non-parametric bootstrapping; central value, lower and upper limits are given. Percent change was estimated using equation (2) for the period described. ^**a**^Island/archipelago ordered North to South. ^**b**^N indicates the most recent count of breeding pairs with the year in parentheses. ^c^Increasing (+); Decreasing (-); Not Determined (ND).

**S2 Table. Database of maximum number of breeding pairs used to estimate the population growth rates for seabirds on the Baja California Pacific Islands**.

**S1 Appendix. Seabird species richness and spatial diversity patterns**.

## Acknowledgments

We would like to thank the following institutions who over the years have supported this research alongside conservation and restoration work for the benefit of the seabirds in the Baja California Pacific Islands: The David and Lucile Packard Foundation, Marisla Foundation, National Fish and Wildlife Foundation, the Montrose Settlements Restoration Program and the S.S. Jacob Luckenbach Trustee Council, Fundación Carlos Slim, Fondo Mexicano para la Conservación de la Naturaleza, the Cornell Laboratory of Ornithology, Global Environment Facility, Programa de las Naciones Unidas para el Desarrollo México, the local fishermen co-operatives and their regional Baja California federation (FEDECOOP), and Comisión Nacional para el Conocimiento y Uso de la Biodiversidad. We are also grateful for the institutional support of the Mexican federal authorities in charge of the islands: Secretaría de Gobernación, Secretaría de Marina, and Secretaría de Medio Ambiente y Recursos Naturales. The work that allowed this research was done under permits from the Unidad de Gobierno-Secretaría de Gobernación, Dirección General de Vida Silvestre-Secretaría de Medio Ambiente y Recursos Naturales, and Direcciones de Áreas Naturales Protegidas-Comisión Nacional de Áreas Naturales Protegidas. This study was partially funded by the Red Temática de Investigación en Áreas Naturales Protegidas (RENANP) of the Consejo Nacional de Ciencia y Tecnología (CONACyT), the CONACyT Basic Science project 251919, and the Centro de Investigaciones Biológicas del Noroeste (CIBNOR). We thank Gabriela Fernández Ham for the graphic design. Finally, we thank all individuals that throughout these years participated in fieldwork phases.

## Author contributions

**Conceptualization:** Federico Méndez-Sánchez, Yuliana-Bedolla-Guzmán, Evaristo Rojas-Mayoral, Alfonso Aguirre Muñoz, Patricia Koleff, Alfredo Ortega-Rubio,.

**Data curation:** Federico Méndez-Sánchez, Yuliana-Bedolla-Guzmán, Evaristo Rojas-Mayoral, Alejandra Fabila-Blanco, Julio Hernández-Montoya, Braulio Rojas-Mayoral, Fernando Solís-Carlos.

**Formal analysis:** Federico Méndez-Sánchez, Yuliana-Bedolla-Guzmán, Evaristo Rojas-Mayoral, Patricia Koleff.

**Funding acquisition:** Federico Méndez-Sánchez, Alfonso Aguirre-Muñoz.

**Investigation:** Federico Méndez-Sánchez, Yuliana Bedolla-Guzmán, Evaristo Rojas-Mayoral, Alejandro Aguilar-Vargas, Alfonso Aguirre-Muñoz, Alicia Aztorga-Ornelas, Esmeralda Bravo-Hernández, Ana Cárdenas-Tapia, Miguel Corrales-Sauceda, Ariana Duarte Canizales, Alejandra Fabila-Blanco, María Félix-Lizárraga, Anely Fernández-Robledo, Julio Hernández-Montoya, Alfonso Hernández-Ríos, Patricia Koleff, Eduardo Íñigo-Elías, Ángel Méndez Rosas Braulio Rojas-Mayoral, Fernando Solís-Carlos.

**Methodology:** Federico Méndez-Sánchez, Yuliana-Bedolla-Guzmán, Evaristo Rojas-Mayoral.

**Project Administration:** Federico Méndez-Sánchez, Yuliana Bedolla-Guzmán, Alfonso Aguirre-Muñoz, Esmeralda Bravo-Hernández, Alejandra Fabila-Blanco, María Félix-Lizárraga, Julio Hernández-Montoya, Fernando Solís-Carlos, Alfredo Ortega-Rubio.

**Resources:** Federico Méndez-Sánchez, Yuliana Bedolla-Guzmán, Alfonso Aguirre-Muñoz.

**Software:** Federico Méndez-Sánchez, Yuliana-Bedolla-Guzmán, Evaristo Rojas-Mayoral, Braulio Rojas-Mayoral, Fernando Álvarez Santana, Maritza Bello Yáñez.

**Supervision:** Federico Méndez-Sánchez, Yuliana Bedolla-Guzmán, Evaristo Rojas-Mayoral, Alfonso Aguirre-Muñoz, Alfredo Ortega-Rubio.

**Validation:** Alfonso Aguirre-Muñoz, Gustavo Arnaud-Franco, Humberto Berlanga-García, Aradit Castellanos-Vera, Eduardo Iñigo-Elías, Patricia Koleff, Alfredo Ortega-Rubio.

**Visualization:** Federico Méndez-Sánchez, Yuliana-Bedolla-Guzmán, Evaristo Rojas-Mayoral.

**Writing – original draft:** Federico Méndez-Sánchez, Yuliana-Bedolla-Guzmán, Evaristo Rojas-Mayoral.

**Writing – review & editing:** Federico Méndez-Sánchez, Yuliana Bedolla-Guzmán, Evaristo Rojas-Mayoral, Alfonso Aguirre-Muñoz, Gustavo Arnaud-Franco, Alicia Aztorga-Ornelas, Humberto Berlanga-García, Esmeralda Bravo-Hernández, Ana Cárdenas-Tapia, Aradit Castellanos-Vera, Ariana Duarte Canizales, Alejandra Fabila-Blanco, María Félix-Lizárraga, Anely Fernández-Robledo, Julio Hernández-Montoya, Alfonso Hernández-Ríos, Eduardo Iñigo-Elías, Ángel Méndez-Rosas, Patricia Koleff, Braulio Rojas-Mayoral, Fernando Solís-Carlos, Alfredo Ortega-Rubio.

